# Prevalence and modulation of rat off-track head-scanning on linear tracks: possible implications for representational and dynamical properties of hippocampal place cells

**DOI:** 10.1101/2025.05.27.656456

**Authors:** Payton J. Davis, Stephen T. Jones, Francesco Savelli

**Author notes:** These authors contributed equally (Payton J. Davis, Stephen T. Jones). Send correspondence to:* Francesco Savelli.

## Abstract

(Re)mapping of different environments by hippocampal place cells is thought to reflect incidental learning. Rat “head scanning” is a spontaneous and presumed investigatory behavior that can trigger the onset of firing locations in place cells. This behavior was studied on (quasi-)circular tracks, and it was speculated that off-track head scans might have been overlooked or inadvertently discouraged in studies employing more common apparatus. To better understand the general prevalence and significance of off-track scanning, we investigated it in rats running laps on linear tracks in rooms featuring visual landmarks. Scanning spanned the length of the track, even in highly familiar conditions and in rats rewarded only at the two ends of the track. Thus, co-localized rewards are not necessary for the occurrence of this behavior. Scanning rate increased markedly in a novel room and then declined steeply during each daily session in this room over 3 days. Transient increases at the beginning of each daily session partially counteracted this decline, producing a “seesaw” profile that is reminiscent of previous observations on place cell plasticity. Therefore, the remapping that place cells are known to undergo in similar contextual changes could conceivably be facilitated by the putative surge of new place fields induced by increased scanning. Investigatory behaviors could thus be causally involved in the representational and dynamic properties of hippocampal representations. Addressing these possibilities offers insight into the incidental creation and update of a cognitive map.

**HIGHLIGHTS:**

- Rat head scanning is known to trigger the onset of firing locations in place cells
- Off-track scanning occurs on linear tracks in familiar and novel conditions
- Co-localized rewards are not necessary for scanning events
- Response to a novel room resembles that seen in place cell dynamics
- Relationships between head scanning and place field formation could help uncover the incidental process of map making

## INTRODUCTION

Hippocampal place cells collectively establish independent maps across different environments, contexts, or behavioral conditions (Bostock et al., 1991; Moita et al., 2004; Leutgeb et al., 2005; Wills et al., 2005; Alme et al., 2014). This (re)mapping process is thought to reflect learning that “is *incremental*…, *latent* because it does not necessarily manifest in behavior, and *incidental* because it takes place in the absence of explicit reward” (O’Keefe, 2007, Ch 11, p. 514, emphasis in original). While examples of behavioral *consequences* of place cell firing remain rare (Lenck-Santini et al., 2002; Gagliardi et al., 2024), it is possible that *spontaneous exploratory and investigatory* behaviors support the presumed incidental creation or update of place cell maps (Lever et al., 2006). The exploration of a novel environment by rodents is characteristically segmented into alternating stops and bouts of forward progression (Golani et al., 1993; Drai et al., 2001; Eilam et al., 2003).

It has long been observed that head movements could reflect internal processing at experimentally defined decision points (Tolman, 1948; Redish, 2016; Kane et al., 2024). Crucially, “head scanning” is also the only documented case of a spontaneous and discrete behavior that can abruptly elicit the onset of new hippocampal place cell firing at arbitrary locations of the environment (Monaco et al., 2014). In the most intriguing display of this phenomenon, a cell fires when the rat samples an *off-track* position by stopping and peering over the track edges—presumably to gather more landmark information. The cell then begins firing consistently in the adjacent location *on the track* in every consecutive lap, thus exhibiting a persistent “place field.” The authors noted how the one-trial character of this phenomenon might reflect the type of neural modifications expected of discrete spatial updates of the map or—more generally—episodic memory function.

Off-track scanning and its impact on place cells and cognition have been characterized only in two studies employing specific types of circular/hexagonal tracks and experimental protocols (Monaco et al., 2014; Rao et al., 2021), leaving questions open about its general prevalence and functional relevance. In particular, it was speculated that this behavior might have been overlooked on linear tracks (e.g., by clipping trajectories exceeding the track edges during data analysis) or inhibited in rats traversing the track ballistically when food reward is restricted to its two ends (Monaco et al., 2014). Still, much of what is known about place fields—from their early characterization (O’Keefe and Speakman, 1987) to their coding properties (McNaughton et al., 1983; O’Keefe and Recce, 1993; Muller et al., 1994), topography (Kjelstrup et al., 2008), statistical distribution (Rich et al., 2014), development (Nitz and McNaughton, 2004), plasticity (Mehta et al., 2000; Frank et al., 2004), and spontaneous remapping (in mice; Zheng et al., 2024)—has been learned over multiple decades in numerous studies employing mazes comprising linear segments that were inspired by the rodent navigation literature (Wijnen et al., 2024). Therefore, it is critical to know whether and how the functional insights gained from the examination of head scanning could retrospectively apply to the existing place cell literature and guide future studies, especially considering that modern approaches often employ virtual reality (VR) strategies based on fixing the animal’s head. Hence, we set out to investigate the expression of off-track head scanning in rats trained to either obtain food reward at the two ends of a linear track (as is common with this apparatus) or throughout the track (as was done in the previous head scanning studies) in both familiar and novel conditions.

## METHODS

### Subjects

Long-Evans rats were obtained from Inotiv Laboratories and allowed to acclimate in the University of Texas at San Antonio Laboratory Animal Resource Center vivarium for at least seven days after arrival (Fig. 1A). Animals were kept on a 12hr dark/12hr light cycle, and all handling, training, and experiments were conducted during the light phase of the cycle (07:00 – 19:00). Female (175 – 199g) and male (200 – 224g) rats were approximately eight and five weeks of age, respectively, at time of arrival to the animal facility. The difference of age was intended to reduce the weight gap between males and females. Animals were balanced with respect to sex (6M, 6F); however, two male rats (Rat 03 and Rat 09) did not complete the experimental protocol (Table 1). Animals were weighed daily. During the training and experimental phases, animals were placed on food restriction to motivate foraging behavior in the experimental apparatus; their weights were maintained at >80% of their ad libitum weight. All experimental protocols were approved by the Institutional Animal Care and Use Committee of the University of Texas at San Antonio.

**Figure 1.**
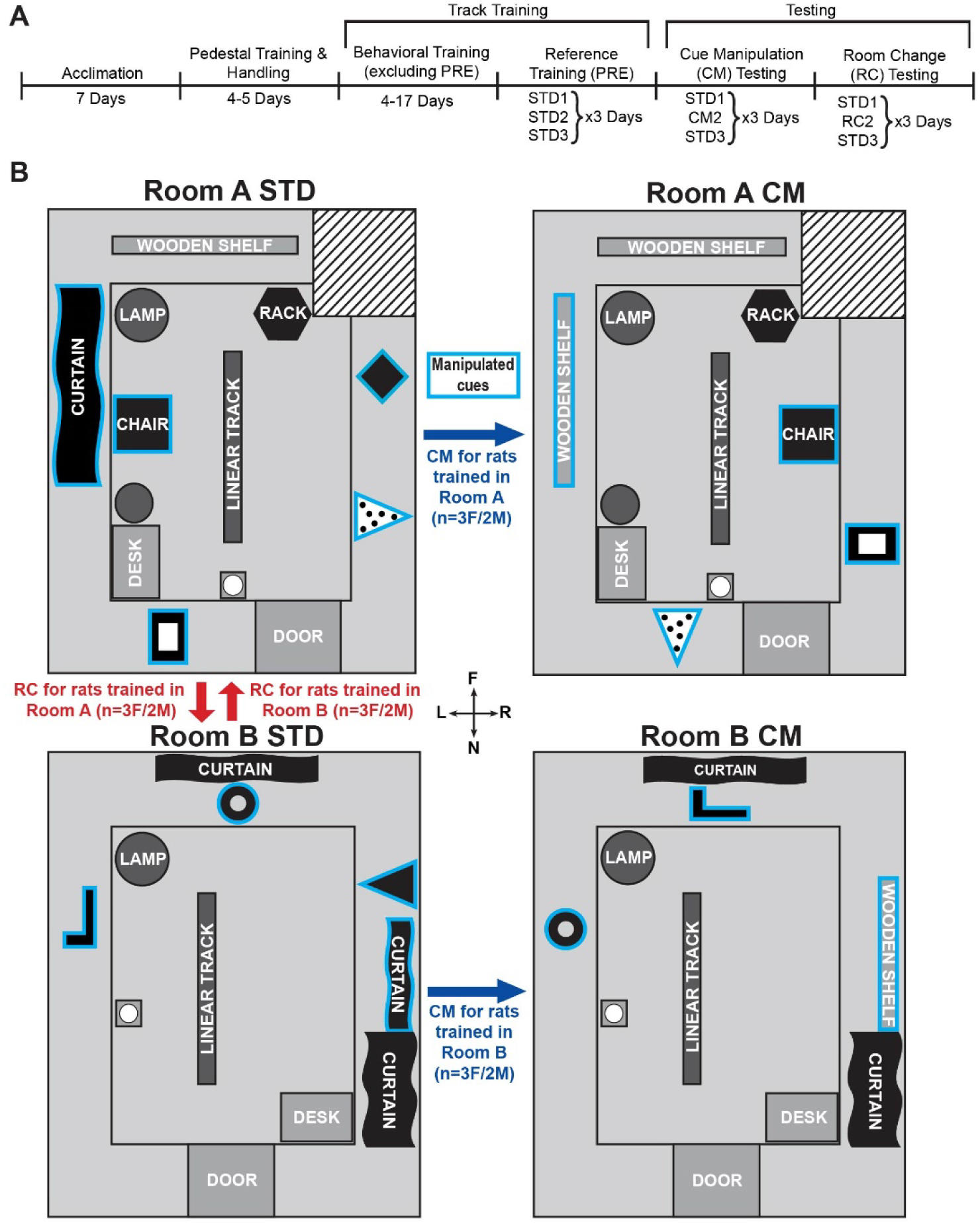
Experimental timeline and manipulations. **(A)** Schematic for the experimental timeline for all rats. The duration of track training varied across animals, based on how quickly they mastered the behavioral tasks. **(B)** Top-down schematics of room layouts and cue constellations for each room and experimental condition. Cues outlined in cyan are manipulated during the cue manipulation (CM) sessions. **(Top left)** Standard (STD) cue configuration for Room A. **(Top right)** CM cue configuration for Room A. The curtain panel is lifted to expose wooden shelves underneath, the chair is moved to the opposite side of the room, the dotted triangle and the rectangular cue are swapped, and the black diamond is removed. **(Bottom left)** STD cue configuration for Room B. **(Bottom right)** CM cue configuration for Room B. Similarly to CM in Room A, a curtain panel is lifted to expose wooden shelves, the black L-shaped and black ring cues are swapped, and the solid black triangular cue is removed. For a given animal, all PRE and CM sessions took place in the room used for training, whereas all RC sessions took place in the other room with its STD configuration. The compass diagram applies to all rooms and configurations: near (N) is the end of the track nearest to the door where the animal begins each session, far (F) is the opposite end of the track, left (L) and right (R) are defined relative to an observer standing at the door and facing into the room.

**Table 1.** Experimental parameters for each rat.

| Rat ID | Sex | Training Room | Reward Location | Track Design | Days of Training (excluding PRE) |
| --- | --- | --- | --- | --- | --- |
| Rat01 | M | Room A | DIST | T-arms | 15 |
| Rat02 | F | Room A | DIST | T-arms | 15 |
| Rat03* | M | Room A | ENDS | T-arms | 16 |
| Rat04 | F | Room A | ENDS | T-arms | 13 |
| Rat05 | M | Room B | DIST | Horse-legs | 12 |
| Rat06 | F | Room B | DIST | Horse-legs | 8 |
| Rat07 | M | Room B | DIST | Horse-legs | 12 |
| Rat08 | F | Room B | DIST | Horse-legs | 7 |
| Rat09* | M | Room B | DIST | Horse-legs | 17 |
| Rat10 | F | Room B | DIST | Horse-legs | 5 |
| Rat11 | M | Room A | ENDS | Horse-legs | 4 |
| Rat12 | F | Room A | ENDS | Horse-legs | 4 |

\*Did not complete full experimental protocol, excluded from analysis

### Apparatus design

Animals were trained on a linear track (250cm L x 12.5cm W, with 2cm lips, elevated 55.5cm from the ground) that followed one of two designs: a black-painted track supported by two grey metal legs that were placed perpendicular to the track at its two ends (“T-arms”, n=4), or a grey-painted track supported by black-painted wooden horse-legs at its two ends (“Horse-legs”, n=8). The track design was modified between the first group of rats (Rats 01-04) using the T-arms version and later animals (Rats 05-12) using the Horse-legs version due to excessive dwelling at the ends of the first track design. Moreover, Rats 01-04 were tested on the same track in all sessions, even when they occurred in different rooms. For all other rats, identically built tracks were used in different rooms, and these tracks were not moved from room to room.

Our experimental design employed two rooms: Room A and Room B (Fig. 1B). Animals were assigned to either room for training and were balanced between the two rooms. Room A was larger than Room B (Room A: 3.05 x 3.76 m with a 0.43 x 0.30 m column in the top-right corner of the room; Fig. 1B top row; Room B: 2.74 x 3.18 m; Fig. 1B bottom row). Room A cues consisted of black curtain panels covering wooden shelves along the walls, a 1.5m tall roll of brown butcher paper standing next to a computer desk, a small black camping chair, a standing lamp, a coat rack covered by a dark table cloth, a black cardboard rhombus wall cue, a rectangular cardboard wall cue with the outer perimeter painted black, and a triangular cardboard wall cue with large black dots (Fig. 1B top row). Room B cues consisted of a computer desk, standing lamp, black curtain panels covering wooden shelves along the walls, and triangular cardboard wall cue that was instead painted fully black. Room B additionally had a black L-shaped wall cue and a black circular ring (Fig. 1B bottom row).

To encourage locomotion, illumination was kept at levels comparable to previous recording studies that found spatial cells responsive to similar wall cues manipulations (Monaco et al., 2014; Savelli et al., 2017).

### Behavioral Training Protocol

After the 7-day acclimation period, animals underwent pedestal and handling training for a minimum of 4 days, wherein they were placed on a circular plate attached to a wooden platform secured by a clamp to a cinderblock (Fig. 1A). The animals remained there for a minimum of 30 minutes, after which they were moved gently onto the laps of laboratory researchers and handled for a minimum of 5 minutes. During the initial handling period, animals were returned to their home cages each day with water and food *ad libitum*. Behavioral training consisted of at least four days of initial exposure to the linear track apparatus in one of the two rooms, Room A or Room B in their standard (STD) cue configuration (Fig. 1B, left column). 25mg chocolate-flavored food pellets from BioServe (Flemington, NJ) were scattered over the length of the track to encourage the rat to explore the track. The rat was removed from its home cage, moved onto the pedestal, and transported into the experiment room. The pedestal was then placed on a cinderblock that remained fixed in the room (Fig. 1B, pedestal location represented by grey square with white circle in all subpanels), and the rat was removed from the pedestal and placed on the end of the track nearest to the door. For the initial exposure period, animals were allowed to explore the track with unrestricted food access (animals consumed an average of 0.5 – 2.5g per session) for at least 20 minutes daily. Animals were placed on food restriction after the first session of the exposure phase of the training process and remained on food restriction until the end of the study.

After the animals had completed four initial exposure days and could consistently traverse the length of the track several times over 20 minutes, they transitioned to the next training phase, in which they started receiving the food reward either exclusively at the two ends of the track (“ENDS” rats) or distributed throughout the main segment of the track (“DIST” rats). In both of these manual reward delivery conditions, approximately five pellets were provided per lap, for a total of 2.5 – 4.5g per session (weight of reward per session was determined by taking the weight of the reward container before and after each session). The experimenter approached the track to deliver reward twice per lap. In some instances, reward was not replenished on a given lap because the animal did not consume the pellets delivered during the previous lap. In this case, we waited for the animal to consume the previous reward before delivering the next. For ENDS animals, the food pellets were placed at each track end as the animal ran to the opposite end of the track (see Video 1). For DIST animals, the experimenter scattered pellets down the track as the animal was running away from where the experimenter was located, to minimize interactions with the animal (see Video 2). The location of the pellets for each lap for DIST animals was thus random and was likely to change between each lap. As the animals continued learning the task, we discouraged doubling back by blocking the animals with a laboratory notebook whenever they began prematurely reversing direction. Excessive grooming and prolonged rests were discouraged by loudly snapping or clapping. All laps were counted using a digital smartwatch or clock that kept track of total elapsed time and lap duration. Once animals were able to complete a set of 30 laps in a single session with smooth and sustained progression (i.e., minimal or no grooming, prolonged rests, or premature attempts to reverse running direction) for a minimum of three training days, they were tasked with running two identical sessions in a single training day. After at least 1-2 days of successful double training sessions, the animals were tasked to complete three consecutive sessions of 30 laps each in a single training day. Animals that completed three consecutive sessions of 30 laps across three consecutive training days were allowed to begin the testing phase (see below), which lasted six days. Between consecutive daily sessions, animals were removed from the training room via pedestal and returned to their home cages with access to water for a minimum of 5 minutes before beginning the next session. During these rest periods and between animals, the tracks were cleaned with 70% EtOH and allowed to air dry before the start of the next session. Animals were not deliberately disoriented between sessions.

All rats from the first cohort (n=4) were trained simultaneously in an open field environment (100cm x 100cm x 50.5cm box, same food type) during a concurrent study. They were trained in the box for 45-60 minutes per day after track training. All animals were retired from box training before the PRE training phase of the present study.

### Testing Protocol

The testing phase consisted of two experiments: cue manipulation (CM) and room change (RC). Each experiment lasted three days, and all animals completed the CM experiment before beginning the RC experiment. Both experimental conditions followed the same general protocol as the training sessions described previously encompassing 3 daily sessions of 30 laps each; however, the second standard (STD) training session was replaced by a CM or RC session (Fig. 1A). In a CM session, the rat was run in the same room where it was trained, after the distal cues were manipulated: two cues were swapped, one cue was removed, and at least one panel of curtains in the room was raised above the shelf it was attached to, therefore exposing wooden shelves that were previously not visually accessible to the rat. These cue-manipulations were room-specific and remained the same for each room for the entire duration of the experiment and for all rats (Fig 1B right column). All cue manipulations between sessions were performed when the rat was resting in its home cage outside of the training/testing rooms. After the CM session, all cues were returned to the familiar, STD configuration so that a third (last) daily session was run in this configuration. Data from this third session were not analyzed in this study. Animals completed three days of CM experiments before they began RC experiments. In the RC protocol, after the animals completed the first STD session in their training room, the second (RC) session was conducted in the opposite room under the STD conditions for that room. For example, an animal trained in Room A would complete a session in Room A with STD Room A cues and then be moved to Room B to run the second session with STD Room B cues (Fig. 1B left column). Animals were not disoriented or otherwise perceptually shielded as they were carried on the pedestal to and from the novel room. All animals were retired from the study and removed from food restriction after completing three days of RC experiments. Data from the last 3 training days constituted the PRE phase we used as baseline in our analyses (Fig. 1A, explained below, in 4/266 sessions included in the analysis, the rat did not finish the 30th lap). For both CM and RC test days, we associate a serial number to each session, reflecting its position in the sequence of sessions run on a given day, irrespective of the type of session. Thus, a typical experiment day will include a sequence of STD1, CM2, and STD3 for the cue manipulation condition or STD1, RC2, and STD3 for the room change condition, while during the PRE phase we had STD1, STD2, STD3.

### Video Collection and Analysis

Video data was captured using an Allied Vision Manta 1.58-megapixel digital camera with an F/1.8 aperture, 1.8-33.0 mm focal length lens (Theia Technologies) mounted ∼2 m above the track. This data was then recorded using the Trodes video capture system (SpikeGadgets, Inc.) and analyzed using DeepLabCut (Mathis et al., 2018). Videos were recorded at 7Hz (to ensure proper camera exposure) or downsampled at this frequency prior to DeepLabCut analysis to ensure a homogeneous treatment of the data.

To apply DeepLabCut-automated tracking to our video data, approximately 15 frames of 1 – 3 videos per animal, comprising 34 videos in Room A and 33 in Room B, were first manually annotated by labeling the ears, nose, and tail base of the animal when visible in each frame. These annotated frames were used for training a resnet50-based neural network, with default parameters. Room A and Room B networks were refined through several rounds of correction-retraining phases to minimize artifacts. We extracted individual example frames that were labeled incorrectly by the model and manually corrected the labels before retraining the network. This process was repeated until automated tracking was deemed satisfactory. Videos from PRE, CM, and RC sessions were then analyzed using the trained networks. We used a likelihood cutoff of 0.5 as the acceptance threshold for each tracked feature. This threshold was applied conjunctively to both ears, with the midpoint used to determine the head position of the animal. Data falling below this threshold was not used (i.e., missing position data in the animal trajectory, see below). Examples of body features automatically tracked via DeepLabCut are shown in Videos 1 and 2.

### Identification and Characterization of Scanning Events

We implemented a custom algorithm to detect laps and off-track scans from position data. The algorithm depended on an accurate delineation of the track’s perimeter within the same video coordinate system used to track the animal’s trajectories. To this end, the vertices and edges of the track were marked and saved as experimental metadata through a feature of the Trodes video acquisition system (SpikeGadgets, Inc.). The coordinates of these features were checked prior to the beginning of the experiments. Offline, the two end zones of the track were defined as the two 25 cm segments at either end. The two track ends were labeled as Near (to the door, “N”, this is where the rats always started training and testing sessions) and Far (“F”, see Fig.1B). We began by parsing the position data obtained through DeepLabCut, filtering out any positions marked as lost or invalid. Laps were defined as trajectories beginning at the N end of the track, moving to the F end, and returning to the N end.

Our scan detection criteria were simplified from those originally developed by Monaco et al. (2014), reflecting differences in apparatus, study design, and our focus on off-track scanning. We defined putative scanning events as instances where the midpoint between the animal’s ears exceeded the bounds of the track by at least 2.5 cm for a minimum of 500 ms (see Videos 1 and 2). For each identified putative scan, we recorded the start and end timestamps and measured the magnitude, or maximum distance from the edge of the track. We categorized scans based on their start position, occurring either at one of the two ends zone or on the main segment of the track. The latter scans were also categorized as occurring on the left or right side of the track. The left and right directions of the experimental room were allocentrically defined relative to an observer standing at the track’s end closer to the doorway and facing the opposite room wall (see labels and compass on Fig. 1B). A final review of these scans was performed to manually remove occasional tracking artifacts showing as outliers inconsistent with the animal’s trajectory as observed in the video.

[[Place Videos 1 and 2 around here]]

### Data Analysis

Data are reported as means and standard errors, and parametric statistical tests (paired and unpaired two-tailed t-tests) were used unless otherwise stated. We employed a one-way MANOVA for our scan duration and magnitude analysis, with experimental condition as a factor. A one-way ANOVA was performed on the data after a significant interaction was revealed by the MANOVA, and significant main effects or interactions were followed up with Posthoc Tukey tests to compare specific differences between groups. A two-way ANOVA with experimental condition and testing room as factors was used to assess differences in lateral scanning biases. Holm-corrected t-tests against zero were used to identify the presence and direction of laterality biases for all groups (statistics reported in Table 2). For the temporal analysis, we also report the 95% confidence interval, defined as two standard errors above and below the mean. We used an alpha level of 0.05 for all statistical tests. Violin plots were generated using Matplotlib’s violinplot function with kernel density estimation to visualize group distributions. The extrema of the data are marked by horizontal bars, with a connecting line indicating the full range. Data analysis and plotting were implemented in Python using open-source libraries (Numpy, Scipy, Pandas, Matplotlib, Statsmodels) made available through the Anaconda Software Distribution (Anaconda, Inc.).

**Table 2.**
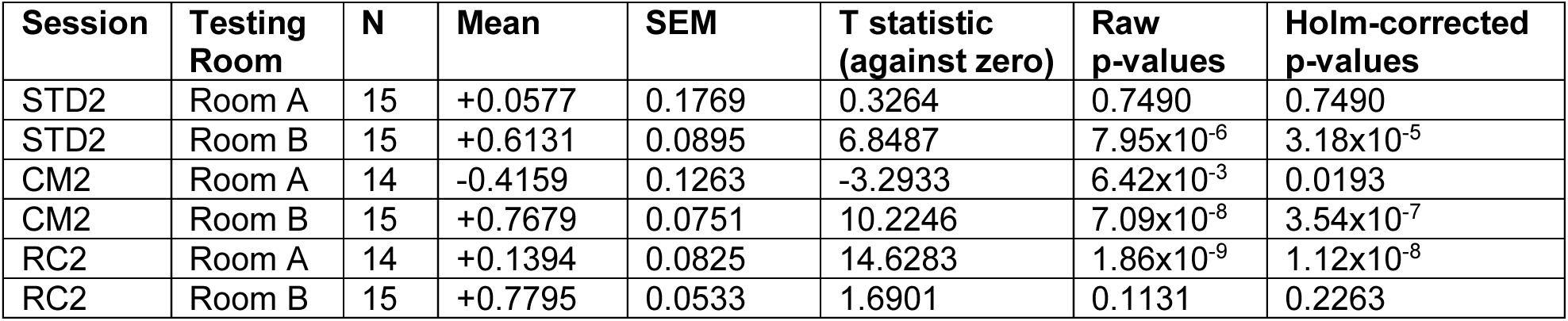
Descriptive statistics and independent t-test results for lateral biases in each condition x testing room condition.

| Session | Testing Room | N | Mean | SEM | T statistic (against zero) | Raw p-values | Holm-corrected p-values |
| --- | --- | --- | --- | --- | --- | --- | --- |
| STD2 | Room A | 15 | +0.0577 | 0.1769 | 0.3264 | 0.7490 | 0.7490 |
| STD2 | Room B | 15 | +0.6131 | 0.0895 | 6.8487 | $7.95 \times 10^{-6}$ | $3.18 \times 10^{-5}$ |
| CM2 | Room A | 14 | -0.4159 | 0.1263 | -3.2933 | $6.42 \times 10^{-3}$ | 0.0193 |
| CM2 | Room B | 15 | +0.7679 | 0.0751 | 10.2246 | $7.09 \times 10^{-8}$ | $3.54 \times 10^{-7}$ |
| RC2 | Room A | 14 | +0.1394 | 0.0825 | 14.6283 | $1.86 \times 10^{-9}$ | $1.12 \times 10^{-8}$ |
| RC2 | Room B | 15 | +0.7795 | 0.0533 | 1.6901 | 0.1131 | 0.2263 |

## RESULTS

We examined *off-track* sampling behavior in rats running laps on linear tracks for food rewards that were either distributed throughout the track (“DIST” rats; n=8) or made available only at its two ends (“ENDS” rats, n=4). We quantified this behavior in three conditions of varying environmental novelty: (1) the familiar room in which training occurred (“PRE”), (2) the same room after several landmarks were removed or shuffled (cue manipulation, “CM”), and (3) the alternate (novel) room that differed in layout and featured novel cues (room change, “RC”) (Fig. 1). One DIST rat and one ENDS rat exhibited inconsistent progress in behavioral performance during training and experiment days and did not complete the full experimental protocol, thus they were excluded from all analyses (see Table 1).

### Scanning in a familiar environment on a linear track

We first asked whether rats would scan off the linear track even in a highly familiar context. We assessed the baseline level of off-track scanning behavior in the training room during the last three training days (PRE days, Fig. 1A), in both training rooms and with both reward delivery conditions. Fig. 2A illustrates examples of PRE trajectories highlighting off-track scanning events. These plots include the scans performed within the two end zones of the track (25cm from either track end, see Methods). Most animals scanned disproportionately in these end zones (64.23% of all scanning events, Fig. 2B, 57.24% in DIST rats and 80.68% in ENDS rats). Hence, for a more insightful comparison to the prior studies using continuous tracks lacking endpoints (Monaco et al., 2014; Rao et al., 2021), we excluded all scanning events occurring within the end zones from all analyses and plots, except when addressing the temporal evolution of scanning rate or whenever examples of whole-session trajectories are illustrated. The remaining scans analyzed in this study occurred anywhere along the rest of the track (Fig. 2B) in both groups of rats (Fig. 2C), averaging about 10 scans per session (mean+-sem: 10.77+-1.00 scans). Note that even the ENDS animals performed scans on this segment of the track away from where they were rewarded (Fig. 2C top). However, in line with considerations put forth by (Monaco et al., 2014), these rats scanned significantly less on this segment than DIST rats (mean+-sem: ENDS, 5.04+-0.68 scans; DIST, 13.22+-1.28 scans; t(88)=-4.05, p=1.08×10^-4^; Fig. 2D). Unlike ENDS rats, DIST rats tended to scan progressively less in locations farther from the N end of the track (from which all rats started each session, see Methods) and closer to F (which was nearer the light source, see Fig. 1B).

**Figure 2.**
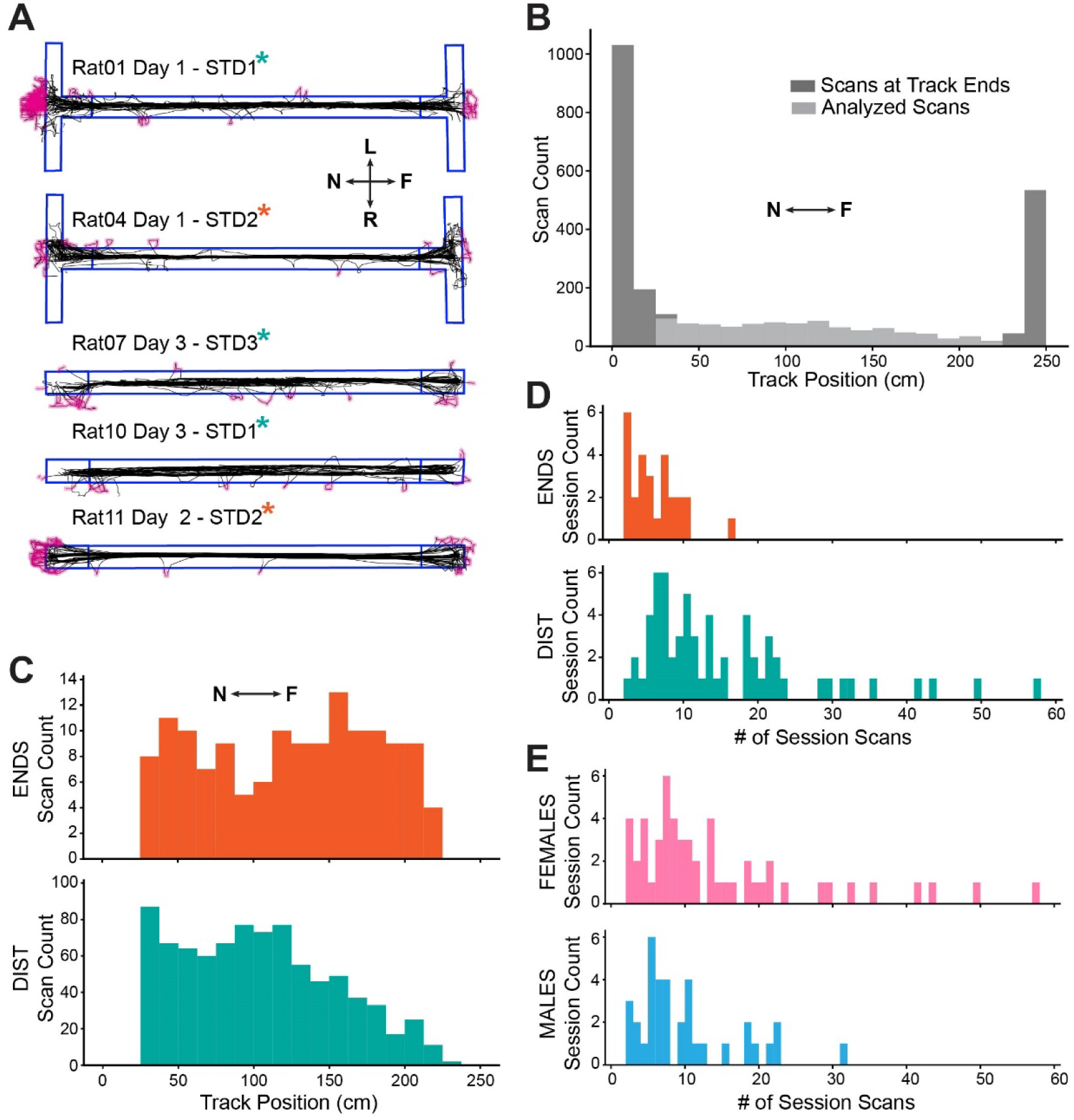
Off-track scanning in the familiar (training) room with different reward delivery protocols. **(A)** Example sessions from the PRE days. Track boundaries are outlined in blue, animal trajectories are shown in black, and scanning events are highlighted in magenta. Rat 01 was trained in Room A and rewarded throughout the track. Rats 04 and 11 were also trained in Room A but were rewarded exclusively at the ends of the track. Rats 07 and 10 were trained in Room B and rewarded throughout the track (see also Table 1). Asterisks denote the reward delivery condition for each rat (same color code as for panels C and D). The compass inset is the same as defined in Fig. 1B. **(B)** Scan counts per 12.5-cm positional bin of the 250 cm track for all PRE STD1, STD2, and STD3 sessions. Dark grey bars represent the scans starting within the end zones of the track (25 cm from each track end; excluded from most analyses; see main text), and light grey bars represent scans starting throughout the rest of the track (the remaining 200 cm between the end zones; included in all analyses). The near end of the track is positioned at 0 cm, the far end at 250 cm (see compass inset).**(C)** The same data represented by the light grey bars in (B) but now separated by reward delivery condition. Animals rewarded at the track ends (ENDS rats) are shown in orange, while animals rewarded throughout the track (DIST rats) are shown in teal. **(D)** Histogram of total scans per session for all PRE STD1, STD2, and STD3 sessions, separated by reward delivery condition. **(E)** The same data as in (D), now separated by sex.

Female rats scanned more than male rats in PRE sessions (mean+-sem: F, 12.46+-1.51 scans; M, 8.22+-0.96 scans; t(88) = 2.11, p = 0.037; Fig. 2E; but see Methods for age differences aimed at reducing weight differences.) Similar sex differences were reported in multiple species for rearing on hindlegs (as discussed in Lever et al., 2006), but another study reported less rearing in females (Sturman et al., 2018).

### Scanning after Cue Manipulation and Room Change

Next, we assessed how the two novelty conditions (CM and RC) presented during the testing phase of the study influenced scanning behavior. We first compared scanning behavior during the first STD session of each CM testing day (STD1) and the immediately following CM session (CM2; the number associated with a session reflects its serial position order in the sequence of daily sessions, e.g., CM2 was the second session run, even if it was the first time the rat was exposed to cue manipulation on that day; Fig. 1A). There was significantly less scanning in CM2 than in STD1 (mean+-sem: STD1, 10.52+-1.48 scans; CM2, 7.86+-1.03 scans; paired samples t-test for all animals and CM days, t(28)=2.70, p=0.012). We reasoned that animals may tend to decrease their scanning over time or distance traveled. If so, they would simply scan less in each subsequent session, and this tendency might counteract any propensity to scan more after the cues were manipulated. To account for this possibility, we first examined how scanning behavior changed between the first and second sessions of the PRE (non-testing) days. Animals indeed scanned less in the second PRE session (STD2; mean+-sem: 8.93+-1.37 scans) compared to the first PRE session (STD1; mean+ sem: 12.53+-1.67 scans; paired samples t-test for all animals and PRE days, t(29)=2.66, p=0.013). Given that this result suggests that animals may scan less during their second session of the day regardless of experimental condition, we attempted to unmask how the novelty of the CM condition could still promote scanning by comparing the changes in scan counts from the 1^st^ to the 2^nd^ daily sessions in PRE vs. CM days (Fig. 3A top and middle). The changes in scan counts did not significantly differ between PRE and CM days (mean+-sem of paired differences STD2-STD1 for all animals and PRE days: −3.60+-1.35 scans; mean+-sem of paired differences CM2-STD1 for all animals and CM days: −2.66+-0.98 scans; unpaired t-test, t(57)=-0.56; p=0.58).

**Figure 3.**
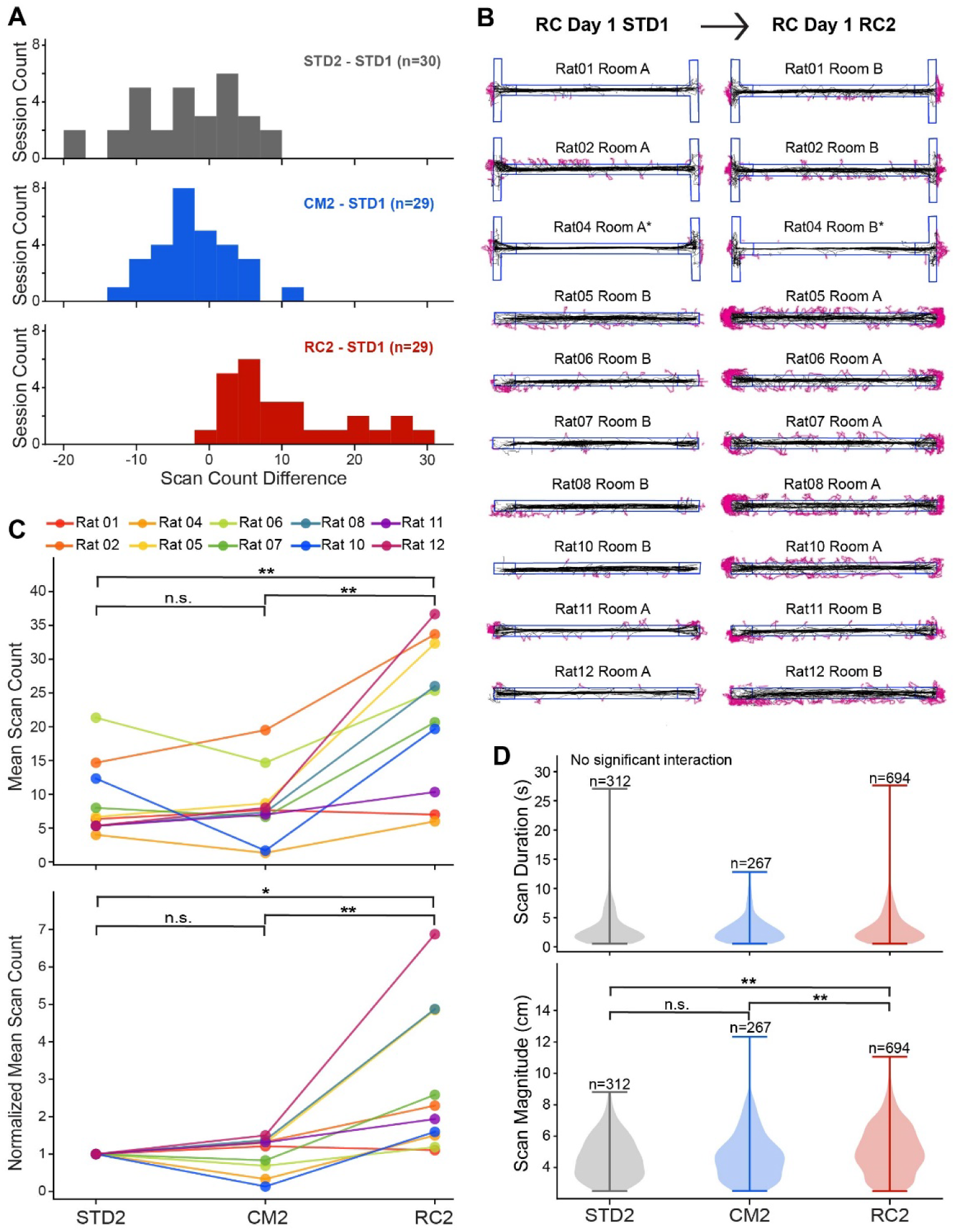
Off-track scanning across novelty conditions. **(A)** Scanning differences in the test sessions for all PRE (familiar; grey), CM (cue manipulation; blue), and RC (room change; red) days. Scan count difference is defined as the difference in total scans between the second (test) and first (familiar) session of a given day (e.g., CM2 total scans - STD1 total scans = CM scan difference). **(B)** Sessions from all 10 rats immediately before and during the first RC experience. Track boundaries, animal trajectories, and scanning events are illustrated as in Fig. 2A. For each rat, the right plot shows the first exposure to the novel room, and the left plot shows the immediately preceding STD session in the familiar room. **(C)** Paired scatter plots of absolute (top) and normalized (bottom) mean scan counts of all STD2, CM2, and RC2 sessions, by rat. In the bottom plot, mean scan counts are normalized to the mean scan count of STD2 sessions for each rat. **(D)** Violin plots of scan duration (top) and scan magnitude (bottom) across STD2, CM2, and RC2 sessions. ***Notes:* \***The sessions from Rat04 in (B) are from the second day of the RC experiment, due to technical issues on the day of recording. n.s. p>0.05; *p<=0.05; **p<=0.01

By contrast, the RC condition elicited an increase in off-track scanning (Fig. 3B). On average, rats performed more than twice as many scans in RC2 sessions as they did in STD1 sessions (mean+-sem; STD1, 9.79+-1.46; RC2, 22.31+-2.29; paired samples t-test for all animals and RC days; t(28)=-6.25, p=9.28×10^-7^). To better appreciate this phenomenon and account for the tendency to decrease scanning between the first and second sessions observed in PRE and CM, we again compared changes in scan counts from the 1^st^ to the 2^nd^ daily sessions across PRE, CM, and RC conditions (Fig. 3A). These changes were greater on RC days (mean+-sem of paired differences RC2-STD1 for all animals and RC days: 12.52+-2.00 scans) than in PRE days (mean+-sem of STD2-STD1 reported above; unpaired t-test, t(57)=-6.71, p=9.59×10^-9^) or CM days (mean +- sem of CM2-STD1 reported above; t(56)=-6.80, p=7.29×10^-9^). Thus, our RC condition strongly promoted increased scanning behavior, unlike our CM condition. We noted, however, that DIST rats entirely accounted for the drop in scan count from STD1 to CM2 (Supplementary Fig. S1A, middle-right panel). ENDS rats, having started from a lower baseline of scan count in STD (as discussed above for PRE days, Fig. 2C-D), responded to the CM manipulation with increased scanning instead (Supplementary Fig. S1A, middle-left panel).

To draw a more direct comparison between STD2, CM2, and RC2 sessions (i.e., just the 2^nd^ daily session across conditions), we computed each rat’s mean total scan count for each condition over the 3 days (Fig. 3C top; averages of the means across rats: STD2, 8.93; CM2, 8.25; RC2, 21.77). Given the considerable variability in individual propensity to scan among rats (see STD2 means in Fig. 3C top and example trajectories in Fig. 3B), we also normalized the CM2 and RC2 means with respect to each rat’s STD2 mean (Fig. 3C bottom; averages of the normalized means across rats: STD2, 1.0 scans; CM2, 1.002; RC2, 2.88). Differences between STD2 and CM2 individual means in Fig. 3C were not significant (paired-samples t-test; absolute values, t(9)=0.45, p=0.66; normalized by STD2 means, t(9)=-0.011, p=0.99). However, when STD2 and RC2 individual means were compared, all animals increased their scanning behavior during RC sessions (paired-samples t-test; absolute values, t(9)=-3.77, p=0.0044; normalized by STD2 means, t(9)=-3.03, p=0.014; Fig. 3C). Additionally, direct comparison between RC2 and CM2 individual means shows that all but one animal (Rat 01) increased their scanning during RC sessions (paired-samples t-test; absolute scans, t(9)=-4.62, p=0.0013; normalized by STD2 means, t(9)=-3.46, p=0.0072; Fig. 3C).

Note that the animals were counterbalanced by training room, to ensure that the impact of the RC condition we observed was due to novelty rather than room-specific features; the difference in scanning between rats tested in different rooms was not statistically significant (absolute means+-sem: Testing Room A RC2, n=5 rats, 24.80+-2.26 scans; Testing Room B RC2, n=5 rats, 18.73+-6.76 scans; t(8)=0.85, p=0.42). The difference in RC scanning by sex did not reach significance in our dataset either (absolute means+-sem: Females RC2, 24.56+-4.47 scans; Males RC2, 17.58+-5.71 scans; t(8)=0.97, p=0.36; see also Supplementary Fig. S1B). The curves of the three ENDS rats in Fig. 3C (Rats 04, 11, and 12) span the variability of the DIST rats; given the small numbers involved and individual differences in the animals’ propensity to engage in scanning, we did not further analyze possible differences between these two groups.

### Duration and magnitude of head scans

Beyond scan counts, we examined differences in the duration and magnitude of scanning events across experimental conditions: STD2, CM2, and RC2. We quantified the magnitudes of scanning events as the maximum distance from the midpoint of the rat’s ears to the track edge (see also Methods). Because scan magnitude and duration were correlated across all groups (Pearson’s correlation, r(1141)=0.40, p=1.15×10^-44^), we performed a MANOVA, revealing a significant effect of experimental condition on duration and magnitude (Pillai’s Trace = 0.020, F(4, 2280)=5.74, p<0.001, ηp²=3.5×10^-5^). Given the significant multivariate effect, we conducted follow-up univariate one-way ANOVAs for scan duration and scan magnitude. Experimental condition did not significantly affect scan duration (F(2, 1140)=2.10, p=0.12, ηp²=0.0037; Fig.3D top). In contrast, experimental condition had a significant effect on scan magnitude (F(2, 1140)=11.42, p=1.2×10^-5^, ηp²=0.020). Post-hoc pairwise comparisons using Tukey’s HSD indicated that scan magnitude was significantly greater during RC2 sessions compared to STD2 (p=0.002) and CM2 sessions (p=0.008), whereas STD2 and CM2 sessions did not differ (p=0.30; Fig 3D bottom).

### Lateral biases

We observed that animals tended to have a directionality bias in their scanning behavior, with more scans occurring on one side of the track than the other (Fig. 4A, additional examples of this phenomenon are in Figs. 2A and 3B). To verify and quantify this observation, we computed a laterality index (LI) for each session by taking the difference between the counts of rightward and leftward scans (relative to the room, see Methods and compass in Fig. 1B) and dividing it by their sum. LI values thus can range from −1 to 1, where negative values indicate a leftward bias, and positive values indicate a rightward bias.

**Figure 4.**
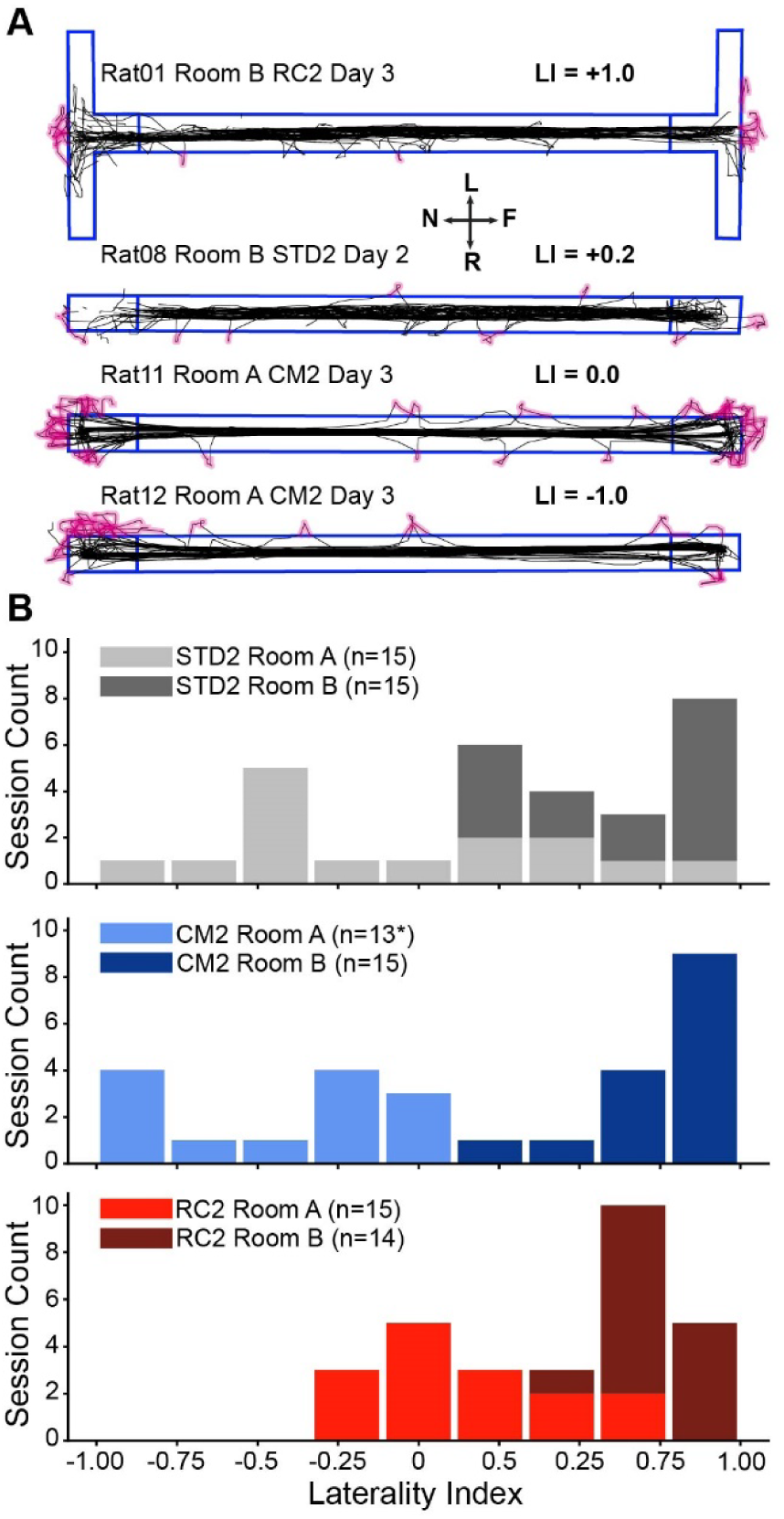
Lateral biases of off-track scanning behavior. **(A)** Example trajectories illustrating varying degrees of lateral bias in scanning, labeled with their laterality index (LI). LIs were computed as the difference between the counts of rightward and leftward scans divided by their sum, such that an LI of −1.0 quantifies a session in which the rat only scanned leftward and an LI of +1.0 quantifies a session in which the rat only scanned rightward. Note that the track masks are accurately positioned relative to the animals’ trajectories, such than any trajectory not aligned with the midline of the track reflects a true bias in the animal’s path (see also Methods; Videos 1 and 2). Example trajectories are illustrated as in Fig. 2A and 3B, including the compass inset as previously introduced in Fig. 1B.**(B)** LIs for STD2 (top; grey), CM2 (middle; blue), and RC2 (bottom; red) sessions, stacked by testing room. In STD2 and CM2 sessions the animals were tested in the same (training) room, whereas the RC data are stacked by the room where the RC session was conducted; for example, the animals that completed their CM sessions in Room A are the same animals represented in RC Room B. ***Note:*** *One session (Rat 04 CM2 Day 3) was excluded from the laterality plots and analyses because the rat scanned only at the ends of the track

We analyzed LI values by a two-way ANOVA with experimental condition (STD2, CM2, RC2) and testing room (Room A, Room B) as factors. There was a strong main effect of testing room (F(1, 247) = 193.83, p < .001, ηp² = .44), indicating a robust difference in scanning laterality biases between testing rooms. The main effect of experimental condition was not significant (F(4, 247) = 1.53, p = .195, ηp² = .024), nor was the experimental condition x testing room interaction (F(4, 247) = 2.17, p = .073, ηp² = .034). Consistent with the effect of testing room on laterality bias, animals tested in Room B exhibited a stronger rightward bias than animals tested in Room A when data were collapsed across experimental conditions (LI, mean+-sem: Testing Room A, −0.091+-0.055; Testing Room B, 0.73+-0.026; t(255)=-13.36, p=1.54×10^-28^). To establish the presence and direction of laterality biases in scanning within each condition x testing room group, we used one-sample t-tests against zero with Holm correction for multiple comparisons (Fig. 4B, Table 2). Animals trained in Room B had a significant rightward bias across both STD2 and CM2 sessions and no clear laterality bias when tested in Room A during RC2 sessions. Animals trained in Room A exhibited no preference in STD2 sessions, a leftward bias in CM2 sessions, and a rightward bias when tested in Room B during RC2 session

Due to the track and cue location in each room (Fig. 1B), these biases can be roughly interpreted as being oriented toward the track side with more room clearance. This was also where the experimenter more often dwelled. So, the scanning lateral biases could be tentatively attributed to a better view of the distal landmarks or greater attention toward the experimenter. Note, however, that while the experimenter may have contributed to orienting scanning behavior, manual reward delivery onto the track cannot have directly triggered most scans or explained their variations across conditions. The experimenter approached the track to deliver reward approximately twice per lap in all sessions regardless of reward delivery and experimental condition (see Methods, and Videos 1 and 2), and scan counts varied by nearly an order of magnitude across sessions in which rats were provided with the same amount of food (Supplementary Fig. S2).

### Evolution of scanning rate over time and distance

Given the rats’ tendency to scan less in consecutive sessions (quantified above), we were interested in the temporal evolution of their propensity to scan within and across days. Thus, we quantified changes in the rate of off-track head scanning over both time and distance traveled, focusing on the second session of the day (STD2, CM2, and RC2). To avoid temporal gaps, we did not exclude scanning events that occurred at the ends of the track from the following analyses and figures.

We first examined scan counts in one-minute time bins (Fig. 5A). Given the high variability in scan counts between individual rats (Fig. 3C), the data in Fig. 5A is normalized relative to each rat’s scan count of the first minute of STD2 Day 1, before averaging across rats. Because session durations, too, vary across rats (Fig. 5B), the number of rats included in the time-bin averages starts decreasing beyond the duration of the fastest session, with the faster rats dropping out first, potentially skewing the interpretation of the scanning rate over time. Therefore, Fig. 5A is limited to the duration of the shortest sessions, focusing on the first six minutes of each session. To overcome these limitations, we assessed how off-track scanning evolved over the distance traveled in the apparatus, as an alternative measure of exposure to environmental novelty on test days. We binned the number of scans occurring over increments of three laps. Again, given the high variability in scan counts across rats (Fig. 3C), we normalized these bins’ counts by the scan count for the first bin (laps 1-3) of STD2 Day 1 for each individual rat, before aggregating the data from all rats, so that each animal has equal weight in the curves plotted in Fig. 5C.

**Figure 5.**
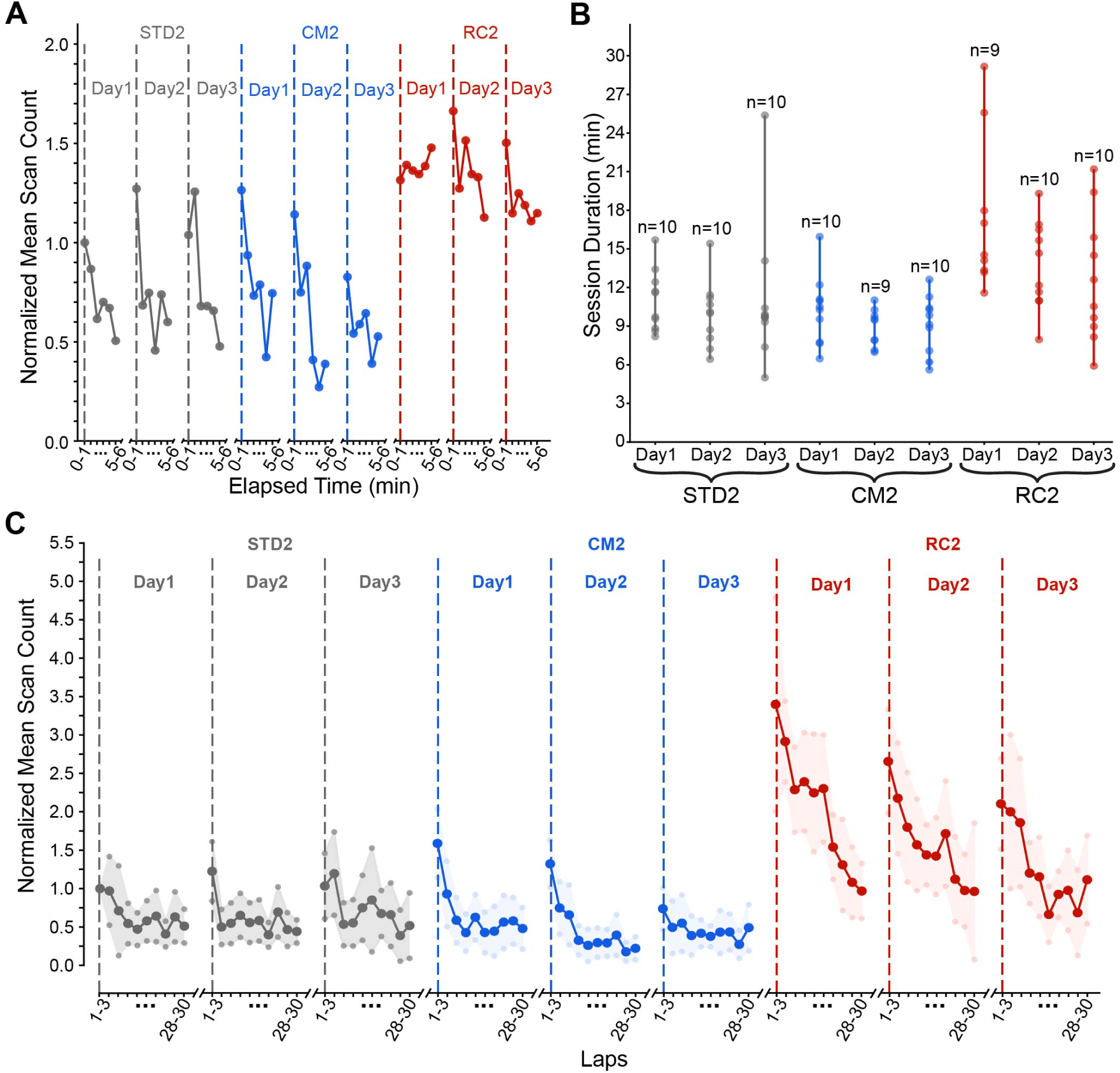
Evolution of off-track scanning behavior. **(A)** Line plot of normalized mean scan counts, binned by minute, up to minute 6 of each session (i.e., truncated to the time the fastest rats took to complete the 30-lap sessions). Before averaging across rats, bin counts were normalized to each rat’s scan count for the first minute of STD2 Day 1. Vertical dashed lines indicate the beginning of each new session. **(B)** Durations of all sessions. Dots represent individual session durations for each rat that completed the corresponding session. **(C)** Line plot of normalized mean scan counts, binned in increments of 3 laps for all 30 laps. Before averaging across rats, bin counts were normalized to each rat’s scan count for the first 3 laps of STD2 Day 1. The shaded region represents the 95% confidence interval (see Methods). Vertical dashed lines indicate the beginning of each new session. ***Notes:*** In these plots and related analyses, scans occurring at the two ends of the track were also included to avoid gaps in the examined timelines.

Intriguingly, the evolution of scanning during CM and RC days followed a declining “seesaw” profile. Animals scanned less during the last laps vs. the initial laps of the session, with scanning gradually declining over the course of the session. However, counteracting resets in scan rate occurred at the beginning of the next day’s corresponding sessions. Because these resets are partial, a slow downward decline over the three days can also be observed in both conditions. These patterns were statistically evaluated through targeted contrasts between the relevant timepoints (e.g., early vs late laps, end of day n vs. start of day n+1, start of day 1 vs. start of day 3, Table 3). All these contrasts were significant on test days except for the overall decline in RC, which yielded a borderline p-value just below conventional statistical significance (p=0.0503). Moreover, these patterns in the evolution of scanning behavior remain evident when the data is replotted separately by reward delivery condition (Supplementary Fig. S3) or sex (Supplementary Fig. S4).

**Table 3.** Summary statistics for paired t-test results for temporal progression of off-track scanning behavior.

| Paired Comparison | N | M1+-SEM | M2+-SEM | T-statistic | p-value |
| --- | --- | --- | --- | --- | --- |
| Laps 1-3 vs. Laps 27-30 (STD2, CM2, RC2/All Days)* | 88 | 1.66+-0.13 | 0.64+-0.08 | 8.23 | $1.71 \times 10^{-12}$ |
| Day 1 Laps 1-3 vs. Day 3 Laps 1-3 (STD2) | 10 | 1.0+-0.0 | 1.03+-0.22 | -0.15 | 0.88 |
| Day 1 Laps 1-3 vs. Day 3 Laps 1-3 (CM2) | 10 | 1.57+-0.23 | 0.74+-0.14 | 4.50 | 0.0015 |
| Day 1 Laps 1-3 vs. Day 3 Laps 1-3 (RC2) | 9 | 3.40+-0.74 | 2.15+-0.34 | 2.30 | 0.0503 |
| Day $n$ Laps 27-20 vs. Days $n+1$ Laps 1-3, with $n=1,2$ (STD2, CM2, RC2)<sup>†</sup> | 29 | 0.59+-0.10 | 1.51+-0.13 | -8.08 | $5.81 \times 10^{-11}$ |

\*Repeated separately for each of the 9 condition-day combination, Holm-corrected $p < 0.026$ for all comparisons except STD2 Day 3, CM2 Day 3, and RC2 Day 3 ( $p > 0.05$ ).
<sup>†</sup>Repeated separately for each of the 6 day-to-day transitions, Holm-corrected $p < 0.017$ for all comparisons except STD2 Day 1 vs. STD2 Day 2, STD2 Day 2 vs. Day 3, and CM2 Day 2 vs. CM2 Day 3 ( $p > 0.05$ ).

## DISCUSSION

Studies of navigation have long appreciated how the task of building a map is inherently circular (Gallistel, 1990; McNaughton et al., 1995, 1996; Etienne et al., 1996; Kuipers, 2000; Thrun et al., 2005). Charting landmarks on a real or cognitive map requires positional estimates of the vantage points from which the landmarks are perceived, hence self-localization—yet self-localization itself presupposes a map. The recursive, probabilistic solutions to this circular problem devised in robotics (Kuipers et al., 2004; Thrun et al., 2005) may be relevant to how the brain makes cognitive maps (Savelli and Knierim, 2019; Fischler-Ruiz et al., 2021; Thurley, 2021). These solutions require active sampling of landmarks integrated with path integration of self-motion cues (McNaughton et al., 2006; Jayakumar et al., 2019). Head scanning, alongside other possibly related behaviors such as rearing on hindlegs, seems to be a behavioral strategy of broad ethological relevance well suited to serve this function (Lever et al., 2006; Monaco et al., 2014; Jun et al., 2016; Rao et al., 2021; Layfield et al., 2023; Shan et al., 2023). The type of head scanning we examined was previously found to reshape the firing patterns of hippocampal place cells under specific behavioral protocols (Monaco et al., 2014). Our observations demonstrate that off-track scanning occurs under experimental conditions and apparatus ubiquitous in place cell studies, even in highly familiar conditions and when reward is spatially restricted. Scans could occur away from reward locations, and overall levels of scanning could vary by up to an order of magnitude even when comparable amounts of reward were delivered, indicating that colocalized reinforcement is not required to trigger this behavior. Intriguingly, scanning surged in a novel room and declined over the course of an experimental session and across days, but with partial daily resets, yielding a dynamic “seesaw” profile reminiscent of novelty-induced changes previously described in place cell plasticity (Frank et al., 2004; Lee et al., 2004; Navratilova et al., 2012). Thus, head scanning offers a compelling yet underexamined window into the incidental formation of cognitive maps and their dynamic properties, linking spontaneous behavior and neural representations.

### Environmental influences

Our CM condition was designed to introduce an intermediate level of novelty between the familiar training condition (STD) and the full room change (RC), in line with the broad place cell literature using partial cue movement, removal, or reconfiguration. Such manipulations tend to produce partial or heterogeneous remapping effects on place fields (e.g., Gothard et al., 1996; Shapiro et al., 1997; Renaudineau et al., 2007), in contrast to the more comprehensive, “global” remapping typically induced by a novel environment (Leutgeb et al., 2005; Wills et al., 2005; Alme et al., 2014). These different responses have been related to memory functions attributed to the hippocampus such as pattern completion/separation (Nakazawa et al., 2002; Guzowski et al., 2004; Leutgeb and Leutgeb, 2007; Colgin et al., 2008; Knierim and Neunuebel, 2016). Our behavioral observations parallel these neural responses in that scanning frequency and magnitude were greater in RC than in CM, and neither parameter generally increased in CM relative to training levels with the standard configuration (PRE). The hippocampus has long been thought to modulate exploratory behavior as a function of perceived novelty (O’Keefe and Nadel, 1978; Lever et al., 2006) and its neural representations have been suggested to reflect latent state inference (Sanders et al., 2020). Accordingly, a manipulation that preserves a coherent global reference frame while altering only a subset of distal cues (as in our CM) may not recruit the same level of exploratory effort as a novel room (as in our RC).

In “double rotation” experiments, Monaco et al. (2014) reported an increase in head scan magnitude as a function of the mismatch angle between proximal (tactile) cues tiling a circular track and a constellation of distal (visual) landmarks. There is no direct analog of the double rotation manipulation for linear tracks. One study recorded place cells on a four-arm radial maze under both double-rotation and cue scrambling/deletion protocols, revealing flexible associations between place fields and subsets of cues (Shapiro et al., 1997). Our CM manipulation differed from the double rotation protocol in several key respects: (i) we did not disorient the rat between sessions; (ii) our cue mismatch only affected distal, visual cues, as opposed to the multimodal combination used in the double rotation; and (iii) we only manipulated a subset of the cues. In contrast, the double rotation was designed to induce an all-encompassing experience (capable of disorienting even the human experimenter, personal observation by F.S.), in which every controllable cue belonged to one of two sets pitted against each other, with no background reference cue intentionally left uncontrolled.

Our use of linear tracks uncovered room-based lateral preferences in head scanning. Head scans were generally biased toward the side of the room with more clearance. Distal/peripheral cues are known to have a stronger influence in setting the orientation of neural correlates of position and head direction in the limbic system (Cressant et al., 1997; Yoganarasimha et al., 2006). Thus, the lateral biases may have been modulated by the perceptual salience of distal cues. The experimenter could also have contributed to orienting the scanning behavior by dwelling more on the side with more clearance, though this factor cannot explain the overall increase of head scans in the novel room. Hippocampal neurons are known to encode aspects of the animal-experimenter interactions (O’Keefe and Dostrovsky, 1971; Snyder et al., 2024). Notably, our experiments were representative of how most place cell studies on linear tracks have been traditionally run, and we actively attempted to minimize these interactions in both reward delivery conditions (see Methods; Videos 1 and 2). While off-track head scanning is prevented in most modern VR apparatus, approaches that avoid head-fixing (Hölscher et al., 2005; Lever et al., 2006; Aronov and Tank, 2014; Grosso et al., 2017; Madhav et al., 2021) could in principle be adapted to study what type of information is sought by the animal during scanning and its effect on place cells (Zeng et al., 2024), for example via closed-loop control of visual cues (Jayakumar et al., 2019; Madhav et al., 2024).

### Relationship with rearing on hindlegs

Parallels between scanning and rearing are worth considering in relation to landmarks. In rearing, the rat rises and stands on its hindlimbs, at times supported by a nearby wall, presumably to gain a better viewpoint on visual cues or access distal olfactory cues (Lever et al., 2006). Rearing seems to be involved in the recognition of distal landmark alterations by pre-weanling rats (Shan et al., 2023). Hippocampal inactivation is most effective at disrupting performance in a spatial task when performed concurrently with rearing episodes (Layfield et al, 2023). Manipulation of cholinergic inputs from the medial septum to hippocampus can alter rearing frequency or duration, possibly reflecting altered attentional states (Monmaur et al., 1997; Cassity et al., 2024).

These considerations make it tempting to speculate that both scanning and rearing could be behavioral manifestations of the same internal drive toward seeking environmental information. Their concurrent investigation is somewhat complicated by the fact that rearing on narrow tracks lacking walls is more precarious and therefore occurs less frequently than in open arenas. (Anecdotally, we observed some rearing, primarily at the two track ends, sometimes blending into off-track scanning, as in some scans in Video 2. However, our experiments were not optimized for rearing detection.) Vice versa, rats routinely engage in rearing in open arenas, but extra-apparatus scanning would only be possible along the perimeters of an elevated platform devoid of walls. Both behaviors are consistent with the intermittent locomotion exhibited by rats exposed to a novel environment (Golani et al., 1993; Drai et al., 2001; Eilam et al., 2003). Future studies could address the possibility of a common neuromodulatory origin (Concetti et al., 2024) and functional overlap between extra/intra apparatus head scanning and rearing. Besides affording better opportunities to acquire distal spatial information, these behaviors could also afford an effective window of integration and plasticity for generating a postsynaptic place field from entorhinal or intrahippocampal inputs (Savelli and Knierim, 2010; Bittner et al., 2015; Davoudi and Foster, 2019; Savelli, 2024). The activity of hippocampal spatial cells correlates with the animal’s head position (possibly the nose; Huxter et al., 2008), and grid cells can anchor their firing patterns to remote visual cues lying beyond the boundaries of the navigable space (Gupta et al., 2014; Savelli et al., 2017). It is therefore conceivable that grid firing patterns could be extrapolated beyond these boundaries during off-track scanning, occasionally aiding in the potentiation of spatially adjacent inputs that would otherwise have remained too weak to generate a place field on the track (Savelli and Knierim, 2010). Indeed, scanning was observed to potentiate place fields in locations where feeble firing was often already present (Monaco et al., 2014), thus unmasking “dormant” place fields, which possibly reflect pre-existing spatial biases inherited from their inputs (McKenzie et al., 2021; Jones and Savelli, 2023; Savelli, 2024; Zheng et al., 2024). Besides grid cells, this process would likely include entorhinal correlates of environmental features possibly attended to by the animal while scanning (Savelli et al., 2008; Solstad et al., 2008; Deshmukh and Knierim, 2011; Høydal et al., 2019; Kinkhabwala et al., 2020). Place cell representations are also known to reactivate (“replay”) during pauses associated with hippocampal sharp-waves/ripples in hippocampal Local Field Potentials (LFPs) and subsequently reorganize across sessions and days (Buzsáki, 2015; Ólafsdóttir et al., 2018; Pfeiffer, 2020). However, the prevalence of theta rhythms during head scanning, together with the abrupt potentiation of place fields associated with this behavior, argues against replay as the proximal mechanism underlying this potentiation. LFP theta power is weaker during scanning than forward running, but stronger in scans than non-scanning pauses (Monaco et al., 2014). The relationship between rearing and theta appears more heterogeneous (Barth et al., 2018).

### Possible implications for place cell properties

The influence of head scanning on the emergence of place fields might extend to their representational properties. For instance, place cells display “global remapping” following context/environment changes similar to our RC condition (discussed above). Conceivably, a surge of new place fields induced by the dramatic increase in off-track scanning that we observed in this condition should contribute to remapping. In environments preventing the animal from peering over the apparatus edges (e.g., in walled boxes), an equivalent influence could be exerted by intra-apparatus scanning (Monaco et al., 2014) or perhaps even rearing (Lever et al., 2006). [Rearing has been shown to modulate place cell firing (Barth et al., 2018), but whether it can induce persistent changes in place fields similar to scanning remains under-investigated.] As a possible illustration of this putative process, place cell remapping was observed even in a familiar, unaltered environment, *after* the rat was subjected to auditory overstimulation in a separate sound chamber (Goble et al., 2009). This manipulation appears to have induced extensive, scan-like trajectory deviations, as compared to the streamlined trajectories that were observed in control and baseline conditions (as suggested by the examples of Fig. 2 vs. Fig. 1 of that study). While this behavior may reflect a delayed anxiety/stress response rather than revived attention to the familiar landmarks per se, it could nonetheless have facilitated the observed place field remapping (e.g., by plasticity acting on new coactivation patterns of position-modulated presynaptic inputs as hypothesized above). In addition to remapping induced by contextual or behavioral changes, place cells are also known to generate distinct maps for either direction of a track, a puzzling phenomenon that seems representationally analogous to global remapping across contexts but could result from large firing rate changes within place fields (Navratilova et al., 2012). As scanning can potentiate barely discernible firing locations into well-defined place fields (Monaco et al., 2014), it could contribute to some of these changes. (Whether scanning-induced potentiation of place fields can proceed independently in the two running directions remains to be investigated, as it has so far been studied only in rats running around a circular track in the same direction.) Together, these considerations suggest a possibly underappreciated role of spontaneous investigatory behaviors in making and differentiating place cell maps. Exploring these possibilities could clarify the nature of learning thought responsible for (re)mapping: namely its incremental, incidental, and latent qualities discussed in the Introduction, especially in light of our demonstration of head scanning’s relative independence from the immediate proximity or overall amount of food reward. An interplay of investigatory behaviors with other internal mechanisms postulated for remapping, such as concurrent changes in grid cell reference frames (Fyhn et al.,2007; Monaco et al., 2011) or autoassociative/attractor neuronal networks (Wills et al., 2005; Colgin et al., 2010), is also possible.

Similarly, the influence of head scanning on the emergence of place fields might also extend to their dynamic properties. After the initial increase in scanning rate upon exposure to a novel room, we observed the rate decline steeply as the session proceeded. This decrease was transiently and partially counteracted by boosts in scanning at the beginning of the first test session each day. These fluctuations give rise to a seesaw-shaped pattern across days that is intriguingly reminiscent of patterns characterizing the plasticity of place fields in novel path configurations (Frank et al., 2004), the emergence of direction sensitivity in place fields (Navratilova et al., 2012; discussed above), and the backward shift of place fields on tracks (which, however, would be more difficult to link mechanistically to head scanning; Mehta and McNaughton, 1997; Mehta et al., 2000; Lee et al., 2004). Once more, it is tempting to draw a parallel with rearing, since preliminary observations suggest that its rate could follow a similar profile in novelty conditions (Lever et al., 2006). Fluctuations in the expression of these investigatory behaviors may thus be causally involved in dynamical aspects of place cell representations. Irrespective of a possible causal link, behavioral and neuronal plasticity still seem bound through shared dynamics induced by novelty.

## Supporting information

Video 1: Consecutive laps, trajectory tracking, and scan events detection in a rat rewarded only at the two track ends.

Video 2: Consecutive laps, trajectory tracking, and scan events detection in a rat rewarded throughout the track.

## AUTHORS CONTRIBUTIONS

**Payton J. Davis:** Designed experiments and analyses; Conducted experiments; Analyzed and interpreted data; Wrote manuscript. **Stephen T. Jones:** Customized behavioral data acquisition and animal trajectory extraction pipelines; Devised/tuned algorithms for lap and scanning detection, feature extraction, and visualization; Assisted with experiments; Gave feedback on analyses and manuscript. **Francesco Savelli:** Conceived study; Designed experiments and analyses; Wrote manuscript; Supervised all aspects of the study.

*CRediT Format.* **Payton J. Davis:** Writing—original draft, Writing—review and editing, Visualization, Validation, Software, Project administration, Methodology, Investigation, Formal analysis, Data Curation. **Stephen T. Jones:** Writing—review and editing, Visualization, Validation, Software, Methodology, Investigation, Data Curation. **Francesco Savelli:** Writing—original draft, Writing—review and editing, Supervision, Methodology, Formal analysis, Conceptualization.

## ACKNOWLEDGMENTS

We thank Roberto Benavidez for apparatus construction; Emily Le, Roberto Benavidez, Priya Raja, Anna Varchuk for assistance with experiments; Kimberly Christian for comments on the manuscript. This study was supported by funding from the University of Texas at San Antonio.

---

**Supplementary Figure S1.**
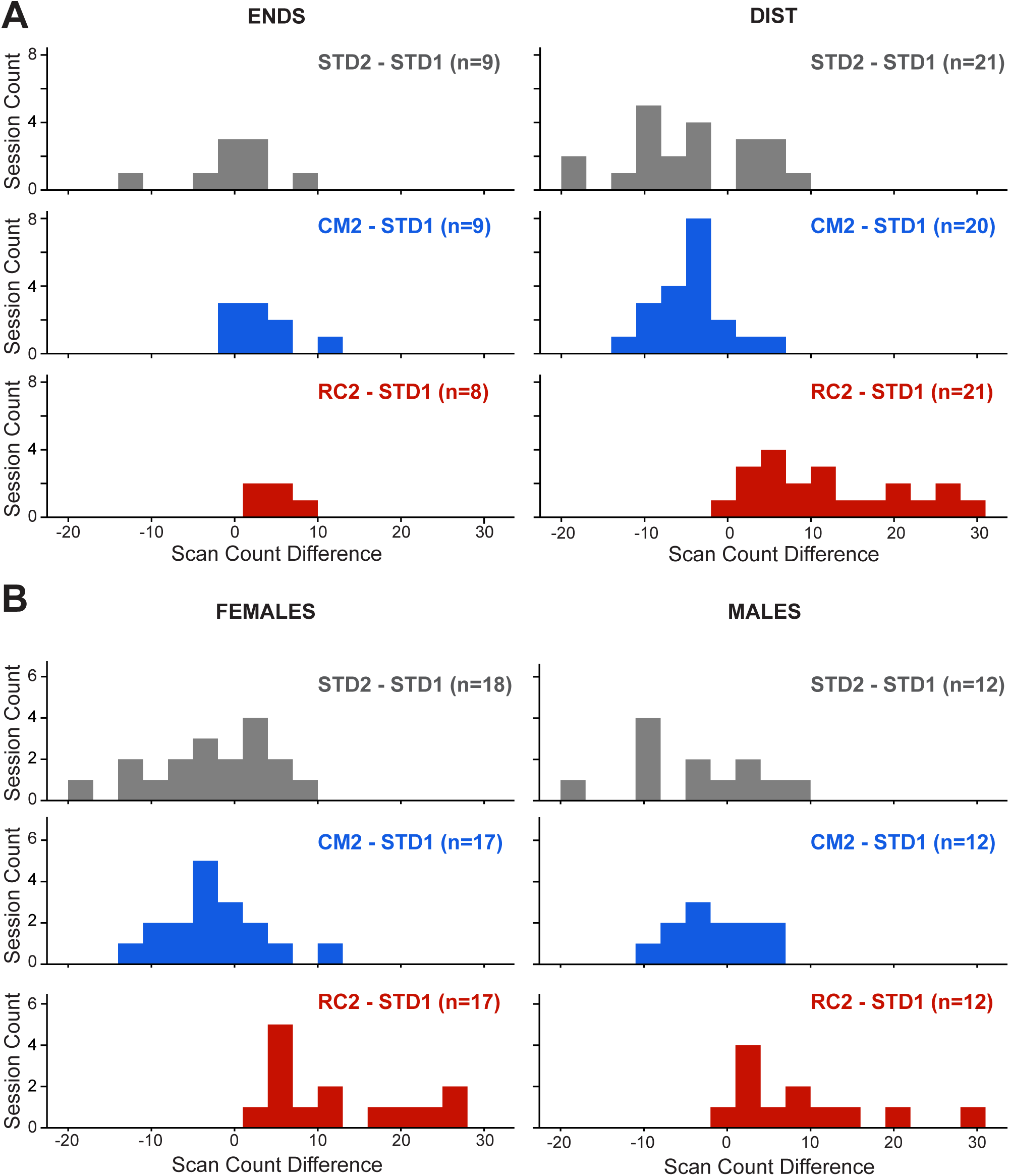
The data shown in Fig. 3A is replotted here, separated by reward delivery condition **(A)** and sex **(B)**.

**Supplementary Figure S2.**
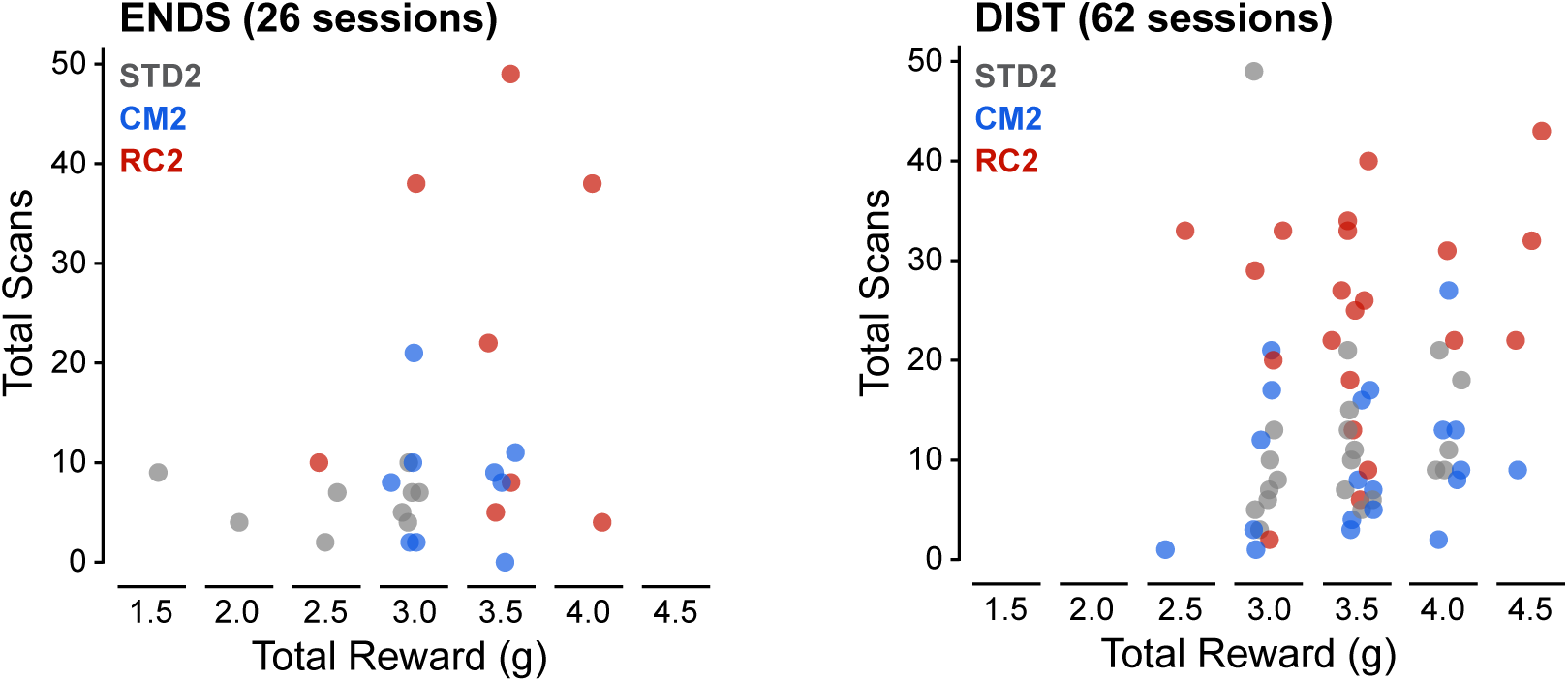
The number of scans can vary by an order of magnitude even across sessions in which a comparable total amount of food reward was provided. Each dot represents a session. Total reward is the cumulative food weight provided over the whole session. Total reward could vary across sessions because, whenever a rat did not consume all or most of the reward on a given lap, the experimenter waited to replenish the reward until the rat consumed it. Total reward was measured with 0.5-g resolution; dots within a given bin are slightly jittered horizontally to improve visibility. (The two sessions in which the animal received <2.5 g of total food reward were both STD2 sessions from Rat04.)

**Supplementary Figure S3.**
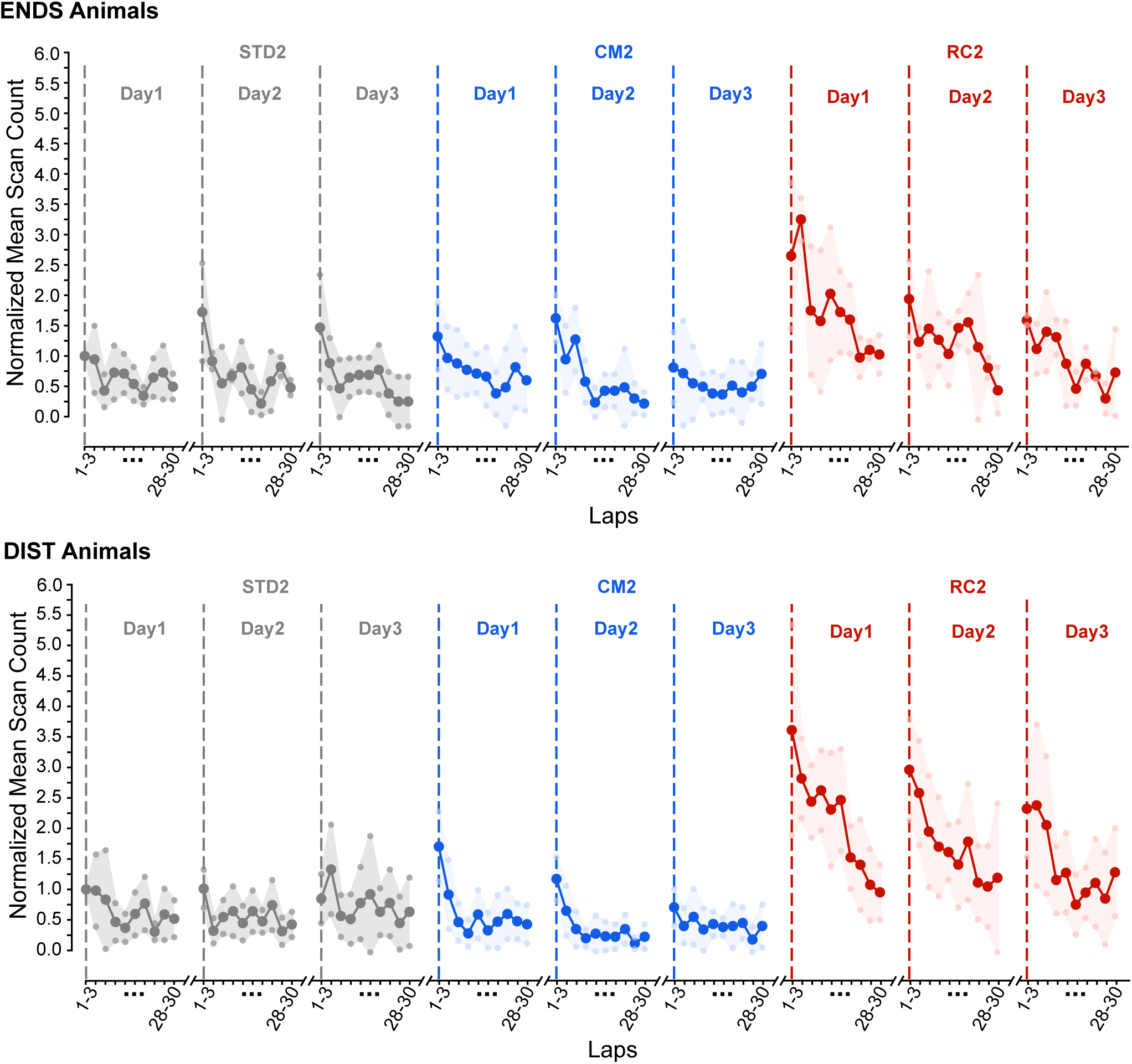
The data shown in Fig. 5C is replotted here, separated by reward delivery condition.

**Supplementary Figure S4.**
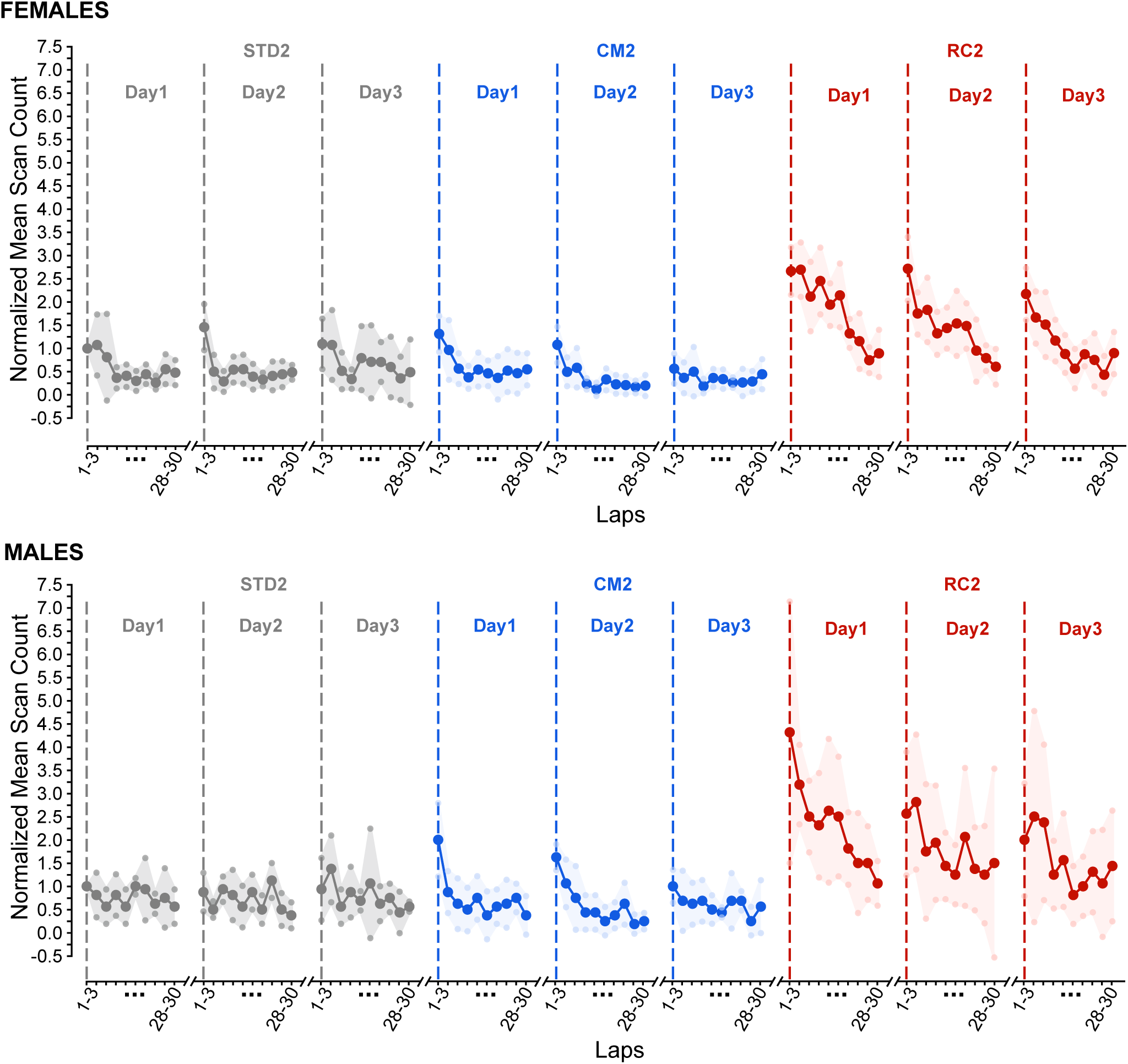
The data shown in Fig. 5C is replotted here, separated by sex.

## Notes

### Competing Interest Statement

The authors have declared no competing interest.

### Summary of Updates

We clarified several methodological aspects of the study and expanded a number of analyses and the discussion. The main revisions include: -Verification of the accuracy of the track masks used to detect off-track scanning events and clarification of the corresponding procedures in the Methods. -Addition of new analyses and figures characterizing the spatial distribution of scanning events along the track, as well as reward-related variables. -Inclusion of two video examples illustrating off-track scanning behavior together with the tracked body features and track mask used in the analyses. -Expanded methodological descriptions regarding food delivery and cue manipulations. -Additional analyses and presentation of results related to sex differences, as well as improved visualization of several data subsets. -Expanded discussion of behavioral and environmental factors that may influence scanning behavior. -Expanded discussion of place cell literature relating to our environmental manipulations.

